# MicroRNA-Mediated Obstruction of Stem-loop Alternative Splicing (MIMOSAS): a global mechanism for the regulation of alternative splicing

**DOI:** 10.1101/2023.04.14.536877

**Authors:** Kai Ruan, German Farinas Perez, Jiaqi Liu, Miroslav Kubat, Ivo Hofacker, Stefan Wuchty, R. Grace Zhai

## Abstract

While RNA secondary structures are critical to regulate alternative splicing of long-range pre-mRNA, the factors that modulate RNA structure and interfere with the recognition of the splice sites are largely unknown. Previously, we identified a small, non-coding microRNA that sufficiently affects stable stem structure formation of *Nmnat* pre-mRNA to regulate the outcomes of alternative splicing. However, the fundamental question remains whether such microRNA-mediated interference with RNA secondary structures is a global molecular mechanism for regulating mRNA splicing. We designed and refined a bioinformatic pipeline to predict candidate microRNAs that potentially interfere with pre-mRNA stem-loop structures, and experimentally verified splicing predictions of three different long-range pre-mRNAs in the *Drosophila* model system. Specifically, we observed that microRNAs can either disrupt or stabilize stem-loop structures to influence splicing outcomes. Our study suggests that MicroRNA-Mediated Obstruction of Stem-loop Alternative Splicing (MIMOSAS) is a novel regulatory mechanism for the transcriptome-wide regulation of alternative splicing, increases the repertoire of microRNA function and further indicates cellular complexity of post-transcriptional regulation.

**One-Sentence Summary:** MicroRNA-Mediated Obstruction of Stem-loop Alternative Splicing (MIMOSAS) is a novel regulatory mechanism for the transcriptome-wide regulation of alternative splicing.

---

Alternative splicing plays an important role in cellular differentiation and stress response in both physiological and pathological conditions. While the vast majority of human genes are alternatively spliced (*1-3*), 22% of disease-causing mutations are splicing sensitive (*4*), and 25% of disease-causing exonic mutations induce exon skipping (*5*). Given that estimated ∼40% of microRNA target sites were found in mRNA coding sequence regions in human cells (*6*), recent evidence points to active roles of noncoding RNAs in alternative splicing. For example, lncRNA can promote the retention of a particular exon in FGFR2 alternative splicing (*7*). Furthermore, double stranded RNAs can affect the splicing of genes associated with muscular dystrophy, while a small nucleolar RNA was found to regulate the splicing of Serotonin Receptor IIC, that is associated with Prader-Willi Syndrome (*8*). Moreover, a study mapping Kaposi’s sarcoma-associated herpes virus (KSHV) microRNA binding sites in B cell lymphoma-derived cells and epithelial cells found that a majority of microRNA target sites were found within the coding sequence of mRNAs, instead of the 3’UTR (*9*). Such observations suggest that microRNA target sites in exonic and intronic regions may point to microRNA’s ability to influence pre-mRNA splicing.

Splicing of long introns is likely facilitated by RNA secondary structure formation in the intron of the pre-mRNA, bringing 5’ and 3’ splice sites into relatively close spatial proximity and promoting excision of a particular intron. While many such long-range interactions are evolutionary conserved and are strongly associated with alternative splicing (*10-12*), a key molecular mechanism is the stem-loop RNA structure formation between complementing sequences, termed “boxes” (**Fig. 1A**). Such box sequences are located near splice sites in introns or exons where the formation of the stem brings distant splice sites together to facilitate long-range splicing (**Fig. 1A**).

**Figure 1.**
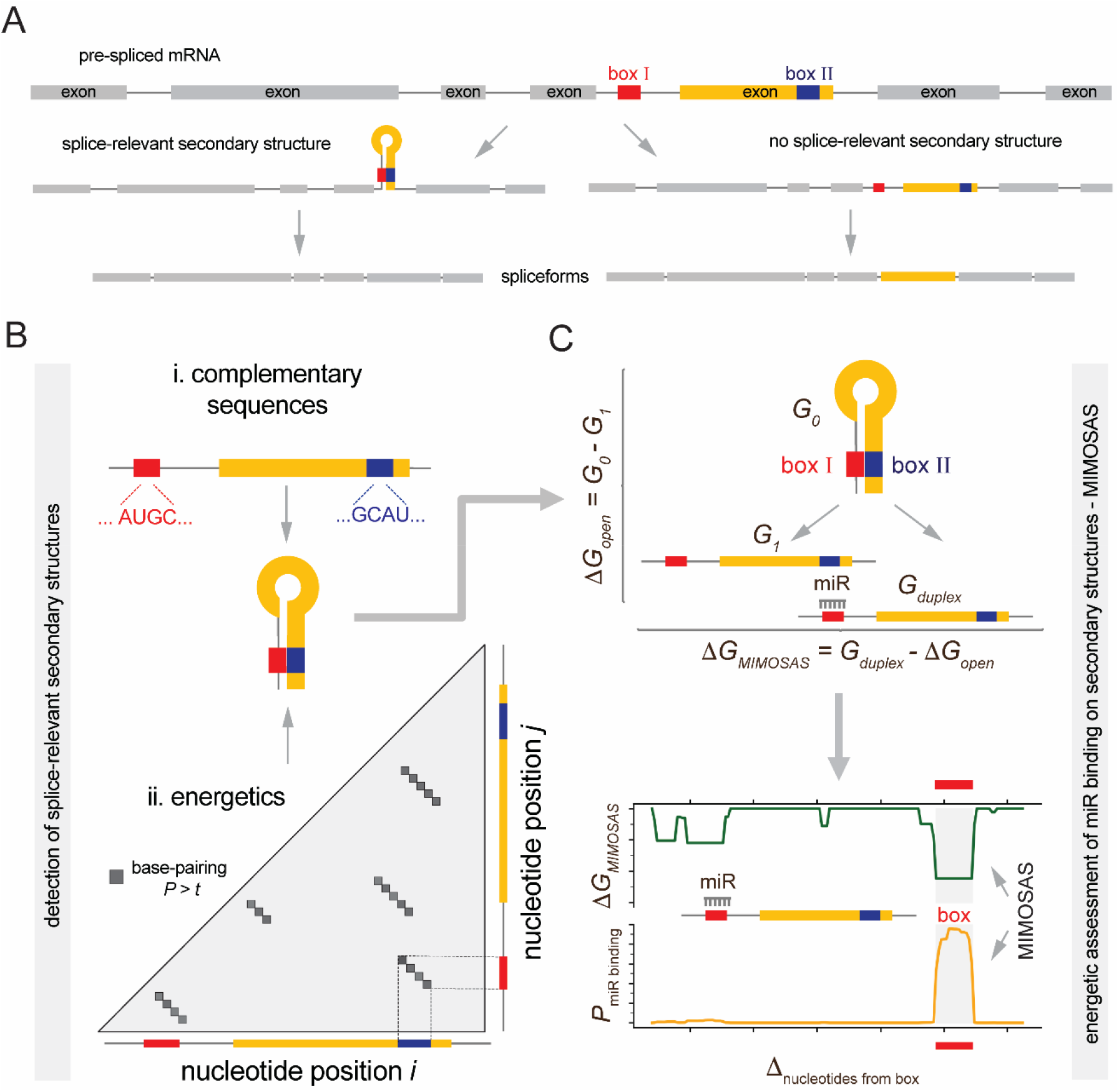
Computational pipeline to detect MicroRNA-Mediated Obstruction of Stem-loop Alternative Splicing (MIMOSAS). (**A**) Our computational pipeline first determines sequences that potentially form secondary structures in mRNA around splice-acceptor and splice-donor sites. (**B**) Such secondary structures are determined through complementary sequences around splice-donor and acceptor sites, termed boxes. Such box candidates are augmented by considering the ensemble of all possible RNA structures, pointing to relevant substructures of mRNAs through probabilities of nucleotide binding. (**C**) Determination of potential binding sites of microRNAs that overlap with boxes are based on the assumption that the energy gained by the microRNA-mRNA binding (*G*_*duplex*_) offsets the energy lost by opening the splice-relevant secondary structures *ΔG*_*open*_ *= G*_*0*_ *– G*_*1*_. If *G*_*duplex*_ *> ΔG*_*open*_ binding of the microRNA that destroys the splice relevant secondary structure is energetically more favorable. Assuming that nascent mRNAs immediately produce secondary structures, predictions are refined through a sequence window upstream a given box, calculating the difference of the duplex with the underlying microRNA and opened structure *ΔG*_*MIMOSAS*_ = *G*_*duplex*_ *-ΔG*_*open*_ as well as the probability that a microRNA binds in the given box region.

MicroRNAs are short noncoding RNAs (∼22nt) that have been well studied in mRNA regulation (*13, 14*), typically by binding complementary nucleotide sequences in the 3’ UTR of target mRNAs through the RNA-induced silencing complex (RISC) (*15*). The vast majority of microRNA-target interactions result in decreased translation of the target (*16*). However, microRNAs have been reported to upregulate translation of their target mRNAs (*17-24*), often through binding the 5’UTR (*19, 22, 23*). Given the prevalence of microRNA-RNA interaction, we predict that the binding of microRNAs can potentially disrupt splice-relevant RNA secondary structures of targets, pointing to an exciting, novel mechanism of splicing and RNA secondary structure regulation (*12, 25*), which we term as MicroRNA-Mediated Obstruction of Stem-loop Alternative Splicing (MIMOSAS). Notably, we have previously identified splice-relevant secondary structures purely on the basis of sequence composition and computational RNA structural modelling of the *NMNAT* pre-mRNA (*26*). In particular, we scanned splice junctions for complementary sequences (i.e., boxes) and corroborated the emergence of a stem-based loop secondary structure that captured intermediate introns (*27*). Based on the assumption that the duplex of the binding microRNA and mRNA is (i) located at the stem of a hairpin loop and is (ii) energetically more favorable than the formation of the stem loop itself, we predicted that miR-1002 potentially interfered with a splice-relevant stem loop formation on the *Nmnat* mRNA that regulated the different splice variants RA and RB. Most notably, we corroborated our computational prediction by experimentally over-expressing miR-1002 that led to the predicted shift of the predicted neuroprotective splice variant RB of the *Nmnat* gene away from RA (*28*).

While our initial consideration was limited to a pair of one microRNA and mRNA, here we show that MIMOSAS is a potentially pan-genomic splicing mechanism. In particular, we designed a computational pipeline that (i) screens for potential sites that form splice-relevant stem-loop RNA structures on different genes, and (ii) computationally predicts microRNAs that disrupt such secondary structures. Most importantly, we experimentally corroborated our computational predictions *in vivo* in *Drosophila* and in cultured human cell models.

## In-silico transcriptome-wide screen for stem-loop mediated alternative splicing

The first step in our bioinformatic pipeline captures the determination of sequences that form secondary structures in a gene’s pre-mRNA around splice-relevant sites (**Fig. 1A**). In particular, we consider complementary sequences around splice-donor and acceptor sites (**Fig. 1B**) by utilizing a recent genome-wide screen in the *Drosophila* genome (*10, 11*). While such an approach points to sequences that are potentially capable of forming a stem, we also considered the energetics of stem formation. In particular, we determined the partition function that captured the entire ensemble of all possible structures (*29*), computing the probabilities that RNA nucleotides indeed bind, and then identified individual structure motifs of high potential splice-relevance (**Fig. 1B**). Such splice-relevant secondary structures are subjected to prediction of potentially overlapping microRNA binding sites, assuming that the energy gained by the microRNA-mRNA binding offsets the energy lost by opening the splice-relevant secondary structures.

Specifically, we determine the energy difference of the mRNA structures with intact (G0) and absent stem loop (*ΔG*_*open*_ = *G*_*0*_ *– G*_*1*_). We consider a microRNA candidate if the energy of the duplex binding structure between a microRNA and mRNA (*G*_*duplex*_) is larger than *ΔG*_*open*_, indicating that the binding of the microRNA that destroys the splice-relevant secondary structure is energetically more favorable (*26, 30*) (**Fig. 1C**). To refine such predictions, we assume that nascent mRNAs immediately produce secondary structures upon emergence from the RNA polymerase. As a consequence, we consider a sequence window upstream of the 5’ end of a box and calculate the energy difference of the duplex with the underlying microRNA (*G*_*duplex*_) and opened structure (*ΔG*_*open*_), *ΔG*_*MIMOSAS*_ = *G*_*duplex*_ *- ΔG*_*open*_, as well as the probability that a microRNA binds a given box (*26, 31*) (**Fig. 1C**).

## In-vivo analysis of microRNAs driving Fas3 splice variants

As testable hypotheses, we first determined splice relevant long-range pre-mRNA secondary structures in the *Drosophila* gene *Fas3*. Utilizing a catalogue of such splice relevant structures that are based on complementary sequences around splice-donor and acceptor sites (*10, 11*), we found a box located in the intron between exon 4 and 5 as well as a box that was located near the 3’ end of exon 5. We refined the prediction of a splice relevant stem by considering the ensemble energy of secondary structures that form on the sequence that captures exon 5 and its 3’ and 5’ flanking introns. In **Fig. S1**, we plotted probabilities that nucleotides interact in the underlying mRNA subsequence, pointing to potential boxes that overlapped with a box that we found through sequence complementarity. Furthermore, we found a stem that captured such boxes (**Fig. S1**) when we determined the energetically most stable structure of the underlying mRNA subsequence. Utilizing these box sequences in **Fig. 2A**, we determined microRNAs that bind box I and II (**Fig. 2B)**. In particular, we sorted microRNAs according to their *ΔG*_*MIMOSAS*_ and chose two microRNAs that target the underlying boxes energetically (un)favorably. We expected that identified candidate microRNAs interfere with the formation of the stem structure, driving the RC splice variant, while we expected splice variants RA/B/D/E/F/G in the absence of MIMOSAS (**Fig. 2C**). To refine our predictions, we considered sequence windows of 200 nucleotides upstream of the 5’ end of a box shifting upstream in increments of the length of the underlying box and determined the energetics of microRNAs binding the mRNA subsequence. Notably, miR-973 showed high *ΔG*_*MIMOSAS*_ and binding probability in the box I region, suggesting that the underlying microRNA is a prime candidate to drive the RC splice variant, while miR-976 does not show remarkable ability to bind in the box region (**Fig. 2D**). In the same way we found that miR-1000 is a prime MIMOSAS candidate binding to box II, while we observed the opposite when we considered miR-999 (**Fig. 2E**). To experimentally corroborate our predictions, we performed real-time PCR after over-expressing corresponding microRNAs in *Drosophila* brains, allowing us to measure the emergence of different splice variants *in vivo*. Indeed, we found that miR-973 and miR-1000 significantly increased the preference of the RC (MIMOSAS) splice variant, while we observed no significant changes with miR-976 or miR-1000 over-expression (**Fig. 2F)**.

**Figure 2.**
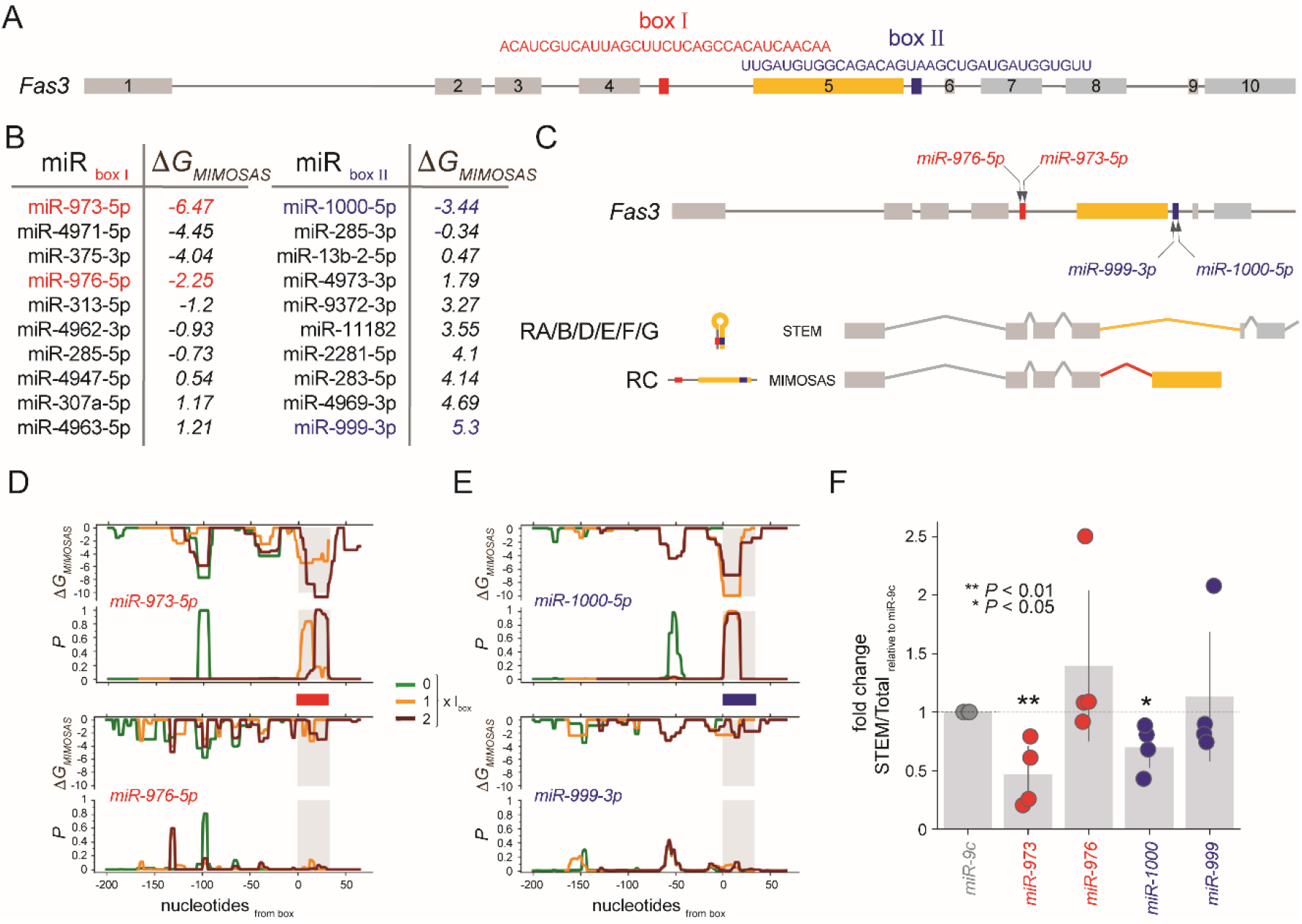
miR-973 and miR-1000 drive splice variants of *Fas3*. (**A**) Structure of the mRNA of *Fas3* indicating box I and II that form a splice-relevant stem around exon 5. Table indicates miRNAs that bind boxes with *ΔG*_*MIMOSAS*_. Energetically (un)favorable microRNAs for further downstream testing are highlighted. (**C**) Splice variant RC that contains exon 5 emerges when highlighted microRNAs thwart stem formation. In the absence of MIMOSAS, *Fas3* splice variants occur that miss exon 5 (RA/B/D/E/F/G). (**D**) *ΔG*_*MIMOSAS*_ and corresponding microRNA binding probabilities of highlighted miR-973 and miR-976 in 200 nt sequence windows upstream of box I. (**E**) *ΔG*_*MIMOSAS*_ and corresponding miRNA binding probabilities of miR-1000 and miR-999 in 200 nt sequence windows upstream of box II. (**F**) The scatter plot of the ratio between the endogenous RA/B/D/E/F/G (STEM) and total mRNA variants measured by qRT-PCR in the brains overexpressing microRNAs by pan-neuronal driver *elav*^*C155*^*-GAL4*. The ratio of STEM vs Total in the miR-9c expressing group was set to 1, and the fold changes were displayed. All data were presented as mean±s.d. ***P≤0.001, **P≤0.01, *P≤0.05, unpaired Student’s *t* test; n=4, triplicate sampling.

## In-vivo analysis of microRNAs driving Nmnat splice variants

Previously, we found that miR-1002 potentially binds and interferes with a splice-relevant stem loop formation on the *Nmnat* pre-mRNA that distinguished between splice variants RA and RB (*28*). Notably, we corroborated our predictions by expressing miR-1002 that led to the expected shift toward the predicted neuroprotective splice variant RB and away from RA. In our previous work, we detected splice-relevant stems in *Nmnat* solely through complimentary subsequences that were located at the splice sites in exon 5 and the upstream intron. To refine the detection of splice-relevant secondary structures, we determined the ensemble of secondary structures of exon 5 and the flanking introns in the pre-mRNA of *Nmnat* (**Fig. S2**) and confirmed that the corresponding stem almost completely overlapped with the boxes we found through complementary sequences. In particular, when the boxes form a stem, the RA splice variant is produced where exon 5 was spliced out. In turn, the RB splice variant emerges when the stem does not form, leading to a retained exon 5 (**Fig. 3A**). Notably, the observed pair of boxes that encapsulate exon 5, can be targeted by numerous candidate microRNAs in addition to miR-1002 (**Fig. 3B**). Specifically, we chose several microRNAs that bind boxes I and II that span a wide range of binding energies. As examples of opposing extremes, we considered miR-137 that binds favorably to box I, while miR-307a as the opposite.

**Figure 3.**
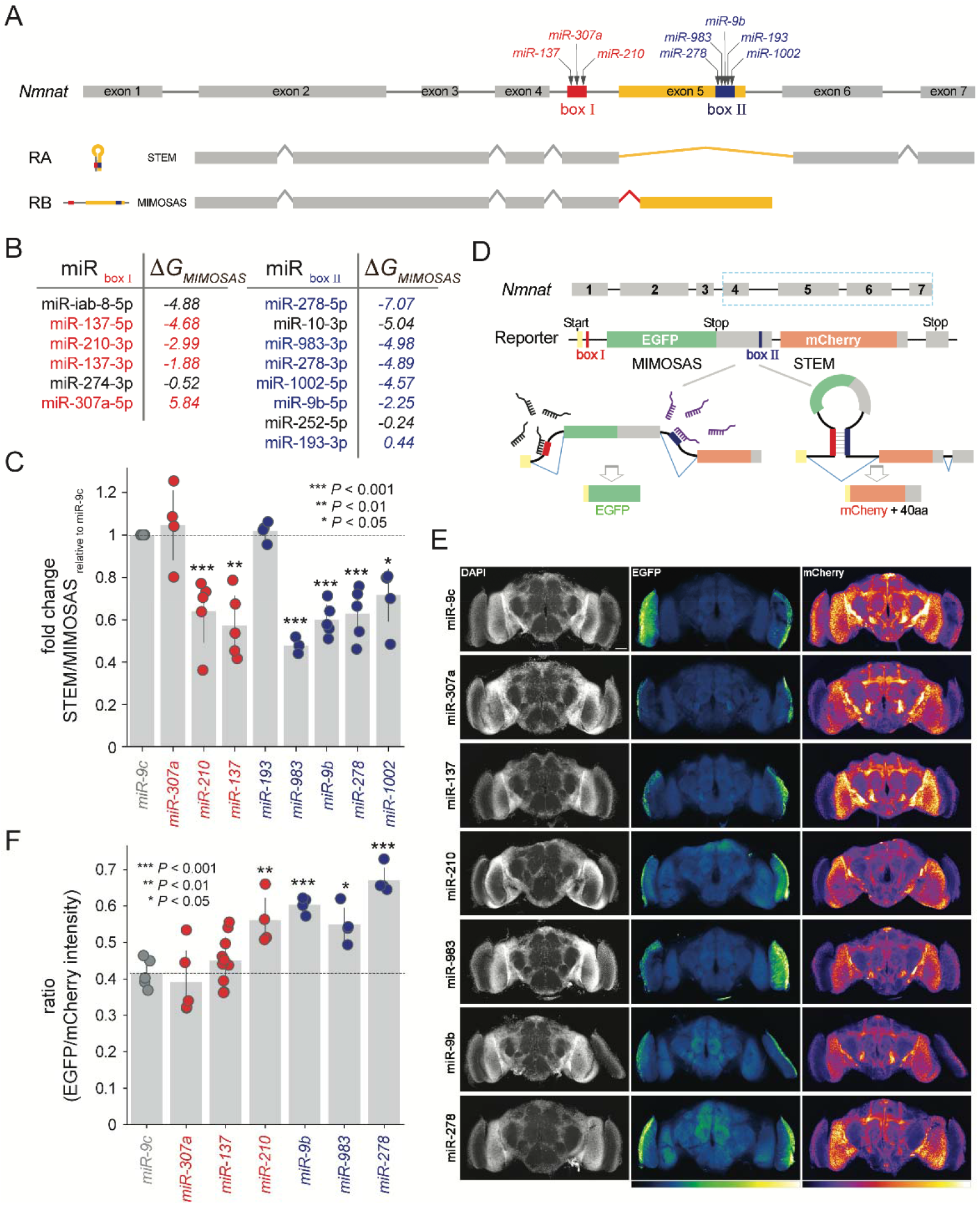
*Nmnat* variants are driven by various microRNAs. (**A**) The gene structure of *Nmnat* harbors a pair of boxes, encapsulating exon 5, that are bound by several microRNAs. If these boxes form a stem, splice variant RA is produced, while MIMOSAS drives the RB splice variant. (**B**) microRNAs that were sorted according to their binding characteristics on boxes I and II. Highlighted microRNAs were chosen for further testing. The scatter plot of the ratio between the endogenous RA (STEM) and RB (MIMOSAS) measured by qRT-PCR in the brains overexpressing microRNAs by pan-neuronal driver *elav*^*C155*^*-GAL4*. The ratio of RA vs RB in the miR-9c expressing group was set to 1, and the fold changes were displayed. All data were presented as mean±s.d. ***P≤0.001, **P≤0.01, *P≤0.05, unpaired Student’s *t* test; n≥3, triplicate sampling. (**D**) An alternative testing method reflects the production of the RB splice variant by EGFP, while mCherry signals the presence of the RA splice variant. (**E**) Brain morphology of 5 days-old (5 DAE) adult flies co-expressing the alternative-splicing reporter and microRNAs by pan-neuronal driver *nysb-GAL4* were imaged by confocal microscopy. The nucleus was marked with DAPI (white) stains, and the intensities of both EGFP and mCherry were indicated with heat maps. The scale bar is 50 μm. (**F**) Quantifying the ratio of EGFP and mCherry intensities in (**E**). All data were presented as mean±s.d. ***P≤0.001,**P≤0.01, *P≤0.05, unpaired Student’s *t* test; n≥4.

Focusing on sequence windows of 200 nucleotides upstream of the 5’ end of the underlying boxes and shifting upstream in increments of the length of the underlying box, we determined the energetics of microRNAs binding such mRNA subsequences (**Fig. S3**). As for miR-137, we observed a strong binding probability with the sequence of box I, while we found no binding signal when we considered miR-307a. Furthermore, we considered miR-278 and miR-983 that showed strong probabilities binding box II (**Fig. S3**). To corroborate our predictions, we performed real-time PCR after expressing corresponding microRNAs or microRNA sponges to reduce endogenous microRNAs. Notably, we found that microRNAs with favorable binding characteristics such as miR-210, miR-137 (targeting box I) as well as miR-983, miR-9b, miR-278 and miR-1002 (box II) drove the emergence of the MIMOSAS-specific RB splice variant. In turn, we found that miR-307a and miR-193 that displayed minimal binding qualities showed no effect in RB production (**Fig. 3C and Fig. S4)**.

To enhance our approach for monitoring splicing events more accurately and to visualize MIMOSAS events directly in live cells, we engineered fluorescence-based genetic reporters to monitor alternative splicing events in a cell type-specific manner *in vivo*. Building upon our previously published alternative splicing reporter for *Drosophila Nmnat* (*32*), we modified the reporter by removing the seven amino acids from the N-terminal of both EGFP and mCherry, which had similar cDNA sequences, including the translational start codon. In particular, we separated EGFP and mCherry by an endogenous intron that contained box I from the *Drosophila Nmnat* gene. Furthermore, we fused the remaining sequence of EGFP with the endogenous 3’UTR of *Nmnat*-RB variant located in exon 5, which was connected by endogenous intron 5, and replaced the DsRed sequence in the original splicing reporter with the remaining sequence of mCherry (**Fig. 3D**). As a consequence, the EGFP fluorescence signal indicates MIMOSAS events, while the mCherry signal points to a STEM facilitated splice product. Notably, our design allows for the accurate measurement of splice variant production, as an incomplete splice product will not have functional fluorescence. To visualize the MIMOSAS events *in vivo*, we generated a transgenic fly line carrying the *UAS-AltSplicing Reporter* and expressed both microRNAs and splicing reporter in the neurons by *nysb-GAL4*. We observed a significant increase in the fluorescent intensity ratio of EGFP (MIMOSAS) and mCherry (STEM) signals in the brains with the co-expression of miR-210, miR-9b, miR-983, and miR-278 compared to the non-targeting (miR-9c) or negative (miR-307a) control (**Fig. 3E-3F**), further confirming our computational predictions (**Fig. 3B**).

## In-vivo analysis of microRNAs driving Rpl3 splice variants

Both *Fas3* and *Nmnat* are examples of alternative splicing that is regulated by a single stem loop secondary structure. To consider more complex secondary structures of pre-mRNAs, we focused on *Rpl3* where we identified nested stem loop structures in addition to the previously reported complementary pair of boxes I and II near splice sites in exon 3 (*10, 11*). By considering the energetically most stable secondary structure of exon 3, we confirmed the formation of a stem between the complementary sequences (**Fig. S5**).

However, when we determined the most stable structure capturing exons 3 and 4, we observed that box II forms a different stem with a newly identified box III in exon 4 (**Fig. S5**). As a consequence of the formation of splice-relevant stems between boxes I and II, or boxes II and III, *Rpl3* pre-mRNA can generate additional splice variants. Specifically, formation of a stem between boxes II/III leads to RA/H production, formation of a stem between boxes I/II leads to RG, and an open confirmation without any stems leads to RD (**Fig. 4A**). Notably, we observed that splice variant RA/H (from stem II/III) is the most abundant, with 94% of total transcripts, while RG (from stem I/II) accounts for 4%, and RD (from no stems open confirmation) accounts for 2% (**Fig. 4A and Fig. S6**).

**Figure 4.**
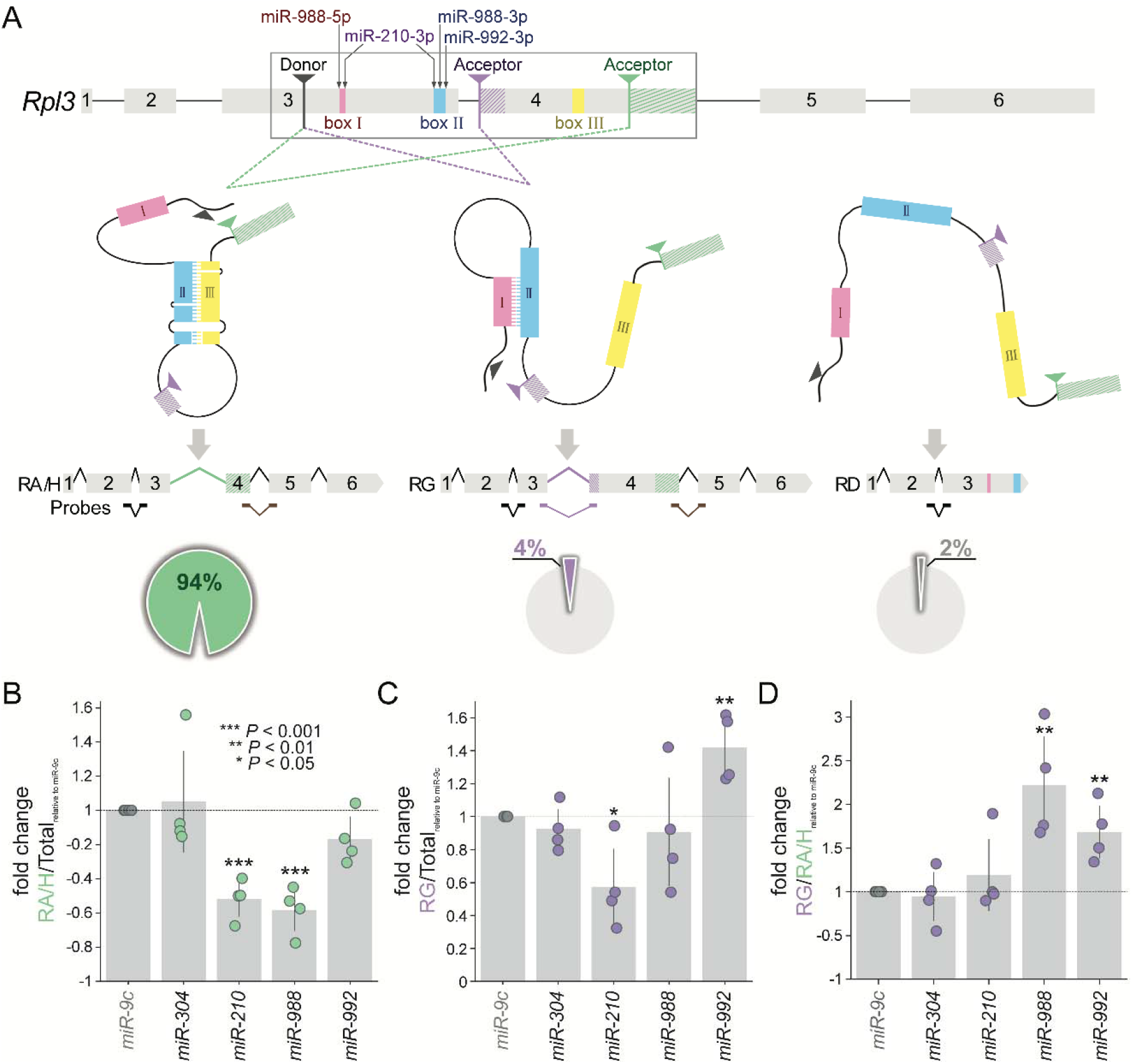
microRNAs drive MIMOSAS on Rpl3 with three competing boxes. (**A**) Diagram of gene structure of RpL3 and predicted spliced mRNA variants. Color coded boxes I, II and III mark the sequences that form the stem-loop structures that are required for alternative splicing. Several microRNAs are indicated to bind box I and II, potentially driving certain splice variants. Black arrow indicates the shared donor splice site. Purple (adjacent) and green (distal) arrows mark the two acceptor splice sites. Three TaqMan real-time PCR probes are marked for the following experiments (**B-D**), the total mRNA variants (black), RA/H and RG (brown), and RG only (purple). The pie diagrams indicate the percentage of endogenous spliced mRNA variants measured by RT-PCR of total RNA from head extracts of adult wild-type flies (yw) (Fig. S6). (**B-D**) The ratio of RpL3 spliced mRNA variants measured by two-color multiplex real-time PCR assay with specific TaqMan probes in flies overexpressing microRNAs. The ratio level in miR-9c overexpressing group was set to 1 and fold change was displayed. All data were presented as mean±s.d. ***P≤0.001, **P≤0.01, *P≤0.05, unpaired Student’s t test; n=4, triplicate sampling.

As a consequence of nested stem loop structures, we expect that the process of stem formation would be highly dynamic with involved box sequences competing for pairing to produce diverse alternative splicing outcomes. Using computational calculations and predictions, we identified several microRNAs that potentially bind to the box regions in *RpL3* and interfere with the stem formation of its pre-mRNA (**Fig. 4A**). Based on both energetically favorable and structurally feasible double criteria, miR-210-3p is predicted to interact with both box I and II, while the 5’ and 3’ arms of miR-988 are predicted to interact with box I and II, respectively. Additionally, miR-992-3p may have a strong interaction with box II (**Fig. S7**). To quantitatively measure the ratio of different *RpL3* mRNA variants after microRNA overexpression, we used a two-color multiplex real-time PCR assay with specific TaqMan probes. In particular, we designed probes for the total mRNA variants, RA/H and RG, and RG only (**Fig. 4A**). Our results showed that expression of miR-210 and miR-988, but not miR-992, significantly reduced the ratio of both RA/H and RG in all four mRNA variants compared to the negative control, miR-304, which had no effect as a non-targeting control (**Fig. 4B**). Furthermore, only miR-210, not miR-988, had the same effect in reducing the portion of RG (**Fig. 4C**). However, miR-992 expression increased the portion of RG among all spliced mRNA variants (**Fig. 4C**). Interestingly, the expression of miR-210 (targeting both boxes I and II) significantly reduced the portions of RA/H (**Fig. 4B**) and RG (**Fig. 4C**) but did not shift the ratio of RG vs RA/H (**Fig. 4D**), suggesting that miR-210 disrupted both stems through MIMOSAS and led to the open confirmation and the splicing of RD. On the other hand, both miR-988 and miR-992 significantly increased the preference for splicing the RG variant (**Fig. 4D**), suggesting that these two microRNAs may disrupt the formation of the stem between boxes II/III and/or enhance the formation of the stem between I/II. While real-time PCR measurements of the endogenous expression of individual variants reveal the effects of microRNAs on splicing and experimentally confirmed the predictions from the computation pipeline, this approach is limited by the availability of variant-specific probes. For example, the expression level of RD can only be inferred or derived from the difference of total transcript and other variants as a consequence of a lack of RD-specific probes.

To directly observe the effects of MIMOSAS on *RpL3* pre-mRNA splicing, we engineered a split-fluorescent proteins (FPs) two color reporter construct that mimics the intricate secondary structures of the pre-mRNA. We retained all the crucial sequences involved in the formation of stems in the pre-mRNA and employed self-complementing split FP (*33*) to monitor the spliced mRNA variants *in vitro*. Utilizing two different FPs, sfGFP and sfCherry, together with nuclear (NLS) and cytoplasmic (NES) localization signals, we distinguished each spliced mRNA variant based on its color and cellular location (**Fig. 5A**). Stem (II/II) favoring the RA/H variant is represented by the expression of cytoplasmic-localized (NES) super fold GFP (sfGFP). In particular, this variant consists of two major parts of split GFP1-10 and GFP11 subunits, a cytoplasmic localization sequence (NES), and a 96bp linker sequence. Such a construct allows two split GFP subunits to form a functional GFP via self-complementation, enabling us to detect green signals in the cytosol. Stem (I/II) that drives the RG variant is represented by the expression of green fluorescent signals in the nuclei. However, when no stems are formed, the RD variant will be transcribed, generating a chimera construct of sfGFP1-10 subunit linked by the sfCherry11 subunit. As a consequence, a red fluorescent signal will be created when combined with the pre-miRNA linked sfCherry 1-10 subunit, indicating a functional sfCherry formed by the complementation of two split subunits. We co-transfected the reporter and pre-microRNA constructs into mammalian COS7 cells for visualization to test the MIMOSAS effect of each candidate microRNA. We selected the cells with red fluorescent signals that indicated the successful co-transfection of two constructs and measured the intensity of each fluorescent channel with confocal microscopy (**Fig. 5B**). Our results demonstrated that miR-210, miR-988, and miR-992 significantly reduced the cytoplasmic GFP intensity (RA/H mRNA variant), compared to the non-target control miR-9c or negative control miR-304 (**Fig. 5C**). Expression of miR-210 reduced the intensity of nuclear GFP signals (RG mRNA variant) (**Fig. 5D**), consistent with real-time PCR results of endogenous RG (**Fig. 4C**). Interestingly, we found that miR-992 expression increased RG (nuclear GFP intensity, **Fig. 5D**) while the intensities of red fluorescent signals grew significantly with both miR-210 and miR-988 overexpression (**Fig. 5E**), confirming our predictions. Collectively, the use of our split-FP dual color reporter allowed us to confirm that miRs-210, 992 and 988 that were predicted to thwart the binding of box II and III indeed diminished the abundance of splice variants RA/H and increased splice variant RD. Most notably, screens of the sequence windows upstream of box I and II suggested that miR-992 does not thwart the formation of a stem, but stabilizes box binding between box I and II, a hypothesis that we experimentally confirmed with increased abundance of splice variant RG.

**Figure 5.**
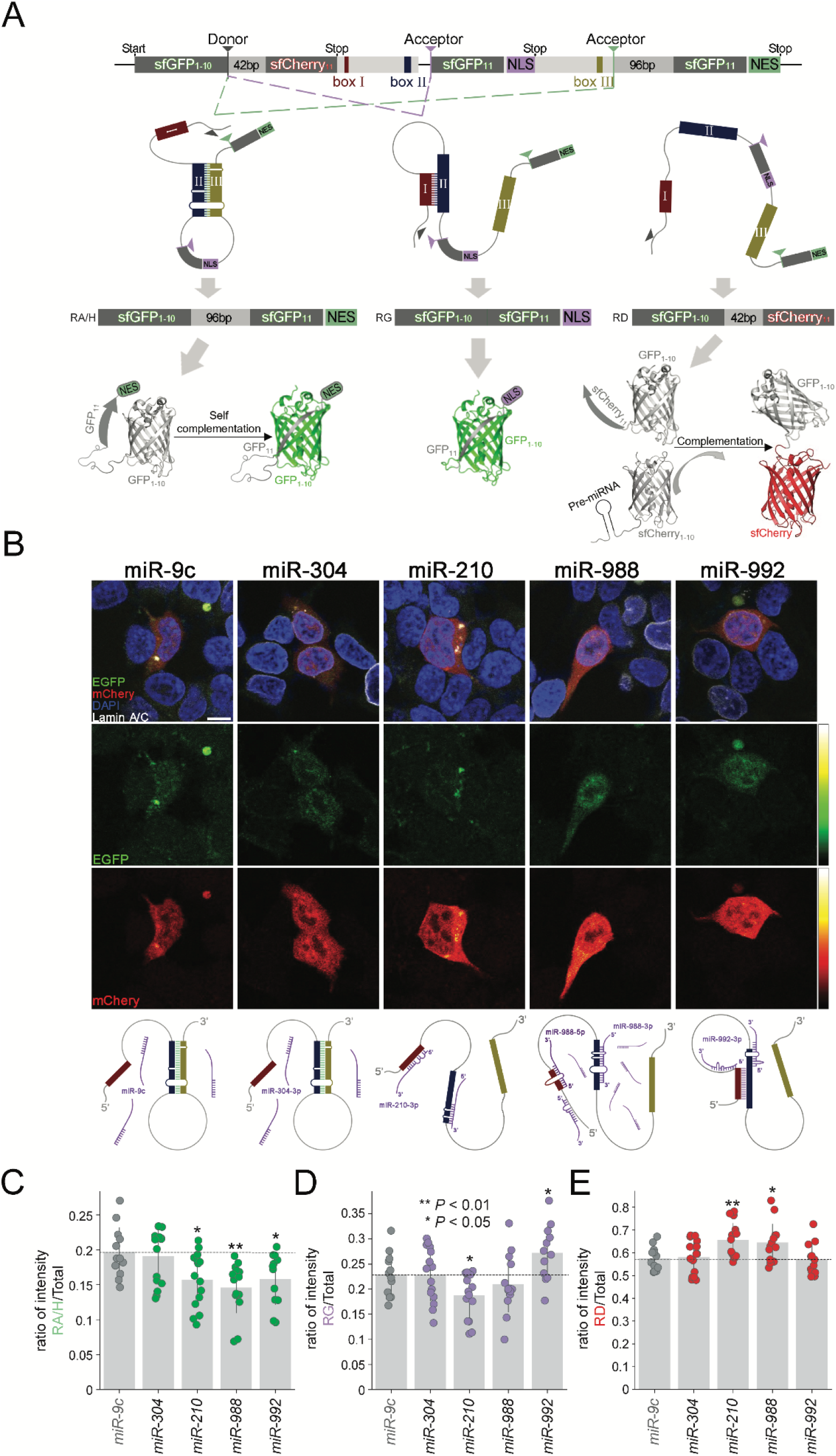
MIMOSAS is driven by microRNAs on the splicing reporter of *Rpl3 in vitro*. **(A**) Design of the splicing reporter for *Rpl3* that includes crucial sequences involved in pre-mRNA STEM formation and self-complementing split fluorescent proteins (FPs), nuclear (NLS) and cytoplasmic (NES) localization signals. Three potential splicing outcomes were accounted for in parallel, each with its spliced mRNA variant and protein product, as well as the visualization mechanism of each reporter. **(B)** Cos-7 cells were transfected with the splicing reporter and sfCherry11-pre-miRNAs (miR-9c, miR-304, miR-210, miR-988, or miR-992), and imaged 48 hours after transfection. The nucleus was marked with DAPI (blue) and Lamin A/C (white) stains, while intensity was indicated with heat maps. Possible schematic diagrams of each microRNA on the splicing reporter are presented below their corresponding column of cell images. (**C**) Quantifying fluorescent intensities, the ratio of spliced variant RA/H is represented by cytoplasmic GFP vs. total fluorescent (GFP + mCherry), **(D)** the ratio of RG is represented by nuclear GFP vs. total fluorescent, as well as **(E)** the ratio of RD is represented by mCherry vs. total fluorescent (all n≥12). Statistical significance through unpaired Student’s *t* test, relative to controls (miR-9).

## Discussion

Although RNA secondary structures have been shown to regulate alternative splicing of long-range pre-mRNA, the driving factors that modulate RNA structure and interfere with the recognition of the splice sites are largely unknown. Essentially, alternative splicing requires the recognition of certain components within a single-stranded pre-mRNA, such as splice sites, branch sites, and cis-acting elements, by trans-acting factors. The formation of RNA secondary structures can inhibit/activate the assembly of spliceosomes through splice site suppression, occlusion/exposure of cis-acting elements, “looping-out” mechanism, as well as the competition between RNA secondary structures (*34*). Long non-coding RNAs (lncRNAs), which are a type of regulatory non-coding RNA, can influence pre-mRNA splicing by altering chromatin, hybridizing with genomic loci or pre-mRNA molecules to create an RNA-DNA or RNA-RNA duplex, or regulating splicing factors (*35*). Conversely, microRNAs, another type of regulatory non-coding RNA, are typically associated with mRNA degradation or translational inhibition, impacting alternative splicing indirectly by post-transcriptionally regulating splicing factors (*36*).

Since majority of splicing events occur co-transcriptionally(*37*), it is likely that microRNA regulate splicing in the nucleus. With the advent of advanced imaging approaches, recent studies have observed the translocation of endogenous mature microRNAs from cytoplasm to nucleus visualized by superquencher molecular probes through optoproation to selectively permeabilize single cells (*38*). Multiple studies have indicated that the in human cells karyopherin XPO1 (Exportin-1) or IOP8 (Importin-8) facilitates the nuclear import of microRNAs-AGO complexes (*39-41*). Even though the detailed function of nuclear microRNAs is not fully understood, currently available evidence suggests that nuclear RISC complexes can modulate the chromatin compaction to achieve a post-transcriptional gene silencing, as well as to regulate the rate of RNA polymerase II procession to affect alternative splicing outcome (*42, 43*). Altering the nuclear level of AGO1, AGO2 or DICER1 has been shown to influence the splicing decision at certain alternatively spliced exons (*28, 42, 44, 45*).

Here, we predict candidate microRNAs that directly interfere and drive alternative splicing through binding with pre-mRNA stem-loop structures. Providing testable hypotheses through a computational pipeline, we experimentally verified the splicing predictions of three different long-range pre-mRNAs in mammalian cell culture models and in vivo in *Drosophila*. Specifically, we observed that microRNAs can either disrupt or stabilize stem-loop structures to influence splicing outcomes, a novel regulatory mechanism for the transcriptome-wide regulation of alternative splicing termed MicroRNA-Mediated Obstruction of Stem-loop Alternative Splicing (MIMOSAS). While microRNAs are mostly studied in the context of mature mRNA 3’UTR binding and target transcript downregulation, MIMOSAS predicts the binding of microRNAs to interior intronic or exonic regions in the pre-mRNA of targets and regulating mRNA splicing. The outcome of such microRNA-pre-mRNA duplex binding allows mRNA expression in an efficient and splice variant-specific manner. Given the dynamic and cell type/tissue-specific expression of microRNAs, MIMOSAS presents a potential new regulatory layer through which cells in different tissues can specifically control the expression of transcript variants. MIMOSAS also unveils regulation of gene expression by noncoding RNAs in a stress or disease state-specific manner and underscores the importance in considering alternative splicing modulation as a therapeutic means. As 60% of point mutations associated with diseases have been found to be related to splicing defects (*46*), several methods have been developed to modify alternative splicing. For example, duplex RNAs have also been used to modify alternative splicing in mammalian cells (*45*), and a method of using single stranded siRNAs has been developed to modify the splicing of a Dystrophin RNA, which is associated with Duchene muscular dystrophy (*47*). Given that such reports indicate the relevance of RNA as splicing modulators our work introduces the possibility of microRNAs as tools to modulate alternative splicing as a potential therapeutic intervention.

## Acknowledgments

We thank Zoraida Diaz-Perez for technical assistance. We also thank Dr John Dezek for critically reading the manuscript.

## Author contributions

Conceptualization: GZ, SW

Methodology: KR, GZ, SW, MK, IH, GP

Investigation: KR, GP, JL, SW

Funding acquisition: GZ, SW

Project administration: SW, GZ

Supervision: GZ, SW

Writing – original draft: GZ, SW

## Data and materials availability

All data, code, and materials used in the analysis must be available in some form to any researcher for purposes of reproducing or extending the analysis. Include a note explaining any restrictions on materials, such as materials transfer agreements (MTAs). Note accession numbers to any data relating to the paper and deposited in a public database; include a brief description of the data set or model with the number. If all data are in the paper and supplementary materials, include the sentence “All data are available in the main text or the supplementary materials.”

## Supplementary Materials

Materials and Methods

Figs. S1 to S7

Tables S1

References (*#1*–*#6*)

