## Supplementary Materials for "MicroRNA-Mediated Obstruction of Stem-loop Alternative Splicing (MIMOSAS): a global mechanism for the regulation of alternative splicing"

**The PDF file includes:**

Materials and Methods

Figures S1 to S7

Tables S1

References (*#1-#6*)

**Materials and Methods**

Plasmid construction

pBID-UASC was a gift from Brian McCabe (Addgene plasmid #35200) (*1*) and pcDNA3.1_sfCherry2 (1-10) was a gift from Bo Huang (Addgene plasmid # 82602 ; http://n2t.net/addgene:82602 ; RRID:Addgene_82602) (*2*). The alternative splicing reporters of *Drosophila Nmnat* and *RpL3* were synthesized and purchased from VectorBuilder. Seven recombinant plasmids were generated for this study. They are pBID-UASC-Nmnat_AltReport v2, pLV[Exp]-puro-EF1A-RpL3_AltReport, and pcDNA3.1_sfCherry2(1-10)-miR-9c (miR-210, miR304, miR-988, miR-992). The primer sequences used in detection and cloning are listed in the Table **S1**.

Fly stocks and culture

Flies were maintained on a cornmeal–molasses–yeast medium at room temperature (22 °C) with 60–65% humidity. All of the *Drosophila* lines were obtained from the Bloomington Stock Center, including *elav-GAL4* and *nysb-GAL4*, as well as the overexpression and knockdown lines of the microRNAs: *UAS-LUC-miRs* and *UAS-mCherry.miRs.sponge.V2*.

RNA Extraction

Total RNA was extracted from at least 40 fly heads per group by FavorPrep tissue total RNA purification kit (Favorgen), according to the manufacturer’s protocol. For each extraction, RNA concentration was measured spectrophotometrically at 260 nm, and 2 µg of RNA was

used for reverse transcription reaction with a high-capacity cDNA reverse transcription kit (Applied Biosystems).

*Drosophila RpL3* Variant Detection

Standard PCR was performed using the FailSafe PCR System (EpiCentre, Chicago, IL, USA) with the following amplification conditions: 25 cycles of 50 seconds at 95°C, 50 seconds at 60°C and 1 min at 72°C. The primers designed (**Table S1**) for variant detection are common forward primer spanning the exon-exon junction between exon 2 and 3, and three distinct reverse primers targeted unique coding sequences for each spliced mRNA variant located in exon 2 or exon 3, respectively (**Fig. S6**).

Real-time PCR

Quantification of mRNA levels was performed using a CFX connect real-time detection system (Bio-Rad) and TaqMan probe-based gene expression analysis (Applied Biosystems). The amplification mix (20 µl) contained 100 ng of ssDNA reverse transcribed from total RNA, and 1 µl of gene-specific TaqMan probe-primer set. The samples were amplified by a two-color multiplex real-time PCR program of 40 cycles of 10 seconds at 95°C, 15 seconds at 55°C, and 1 min at 72°C. The quantification of mRNA levels was carried out by the 2(-Delta Delta C(T)) Method (*3*). Seven TaqMan probes were used in this study. They are FAM-Dm02150883_g1, VIC-Dm02144515_g1, FAM-Dm01810909_g1, VIC-Dm01810910_m1, FAM-Dm02135667_g1, FAM-Dm02135669_g1, VIC-Dm02148683_g1.

In-cell reporter assay

Cos-7 cells were co-transfected with lipofectamine 2000 (Life Technologies). 1-2 µg of cDNA was diluted into 100 µl of Opti-MEM I Medium (Invitrogen) and mixed gently. Lipofectamine 2000 mixture was prepared by diluting 2-4 µl of Lipofectamine 2000 in 100 µl of Opti-MEM I Medium. The ratio of DNA to Lipofectamine 2000 used for transfection was 1∶2 as indicated in the manual. The DNA-Lipofectamine 2000 mixture was mixed gently and incubated for 20 min at room temperature. Cells were directly added to the 200 µl of DNA-Lipofectamine 2000 mixture. After 48 h transfection, cells were fixed and stained with 4,6-diamidino-2-phenylindole (DAPI) and Lamin A/C conjugated Alexa Fluor^®^ 647 (1:500, Cell Signaling Technology #41357). Samples were visualized with an Olympus IX81 confocal microscope under × 60 magnification.

In-Fly reporter assay

Adult brains were fixed in phosphate buffered saline (PBS) with 3.7% formaldehyde for 15 min and washed in phosphate buffered saline with 0.4% Triton X-100. DAPI (1:1000, Invitrogen) staining was performed after 3 times wash. Before imaging, tissues were mounted on microscope slides in Vectashield Mounting Medium for Fluorescence (Vector Laboratories).

Confocal Image Acquisition and Processing

Confocal microscopy was performed with an Olympus IX81 confocal microscope and processed using FluoView 10-ASW (Olympus) or ImageJ (NIH), and Adobe Photoshop 2023 (Adobe, USA).

Quantification and Statistical Analysis

Data were represented as mean ± s.d.. Statistical analysis was performed in GraphPad Prism using unpaired Student’s *t*-test. Values of P < 0.05 were considered statistically significant.

Selection of candidate stems

Starting point for the candidate selection was a list of evolutionary conserved complementary regions, termed boxes, identified in (*4*). For each pair of boxes in a gene of interest, we checked whether the secondary structure induced by pairing of the boxes could influence isoform selection. Minimum free energy (MFE) secondary structures and base pairing probabilities were computed using the RNAfold program of the ViennaRNA package (*5*) for a region of the pre-mRNA encompassing both boxes and nearby exon-intron boundaries to ensure that the boxes indeed from a stem with high probability. In some cases, additional pairs of boxes were identified in this step.

Identification of MIMOSA miRNAs with RNAup

For each miRNA with a potential binding site overlapping our boxes, the secondary structure that would be formed after miR binding, was predicted using RNAfold with constraints that forbid the target site to form intra-molecular base pairs. The predicted structures were inspected to check whether the box interaction was indeed destroyed as well as to compute the opening energy for miR binding as the difference between the folgin energies with and without constraints. Moreover, we used RNAup (*6*) to compute the strength of interaction between the miRNA and mRNA. The free energy of binding computed by RNAup consists of the duplex energy, i.e. the free energy of the intermolecular duplex formed between miR and mRNA, as well as the opening energy necessary to make the binding sites accessible for interaction, *ΔG_MIMOSAS_* = *G_duplex_ – ΔG_open_.* In principle, the binding free energy allows to compute the fraction of bound mRNAs if both miR and mRNA concentrations were known. In addition, RNAup computes the conditional probability that the interaction covers a given position. A value close to 1 ensures that no other favorable binding sites exist in the region considered. Since most splicing events occur co-transcriptionally, we mimicked the effects of co-transcriptional folding by performing RNAup computations for three different windows along the mRNA, roughly representing three different time points during transcription.

**Supplementary Figures**

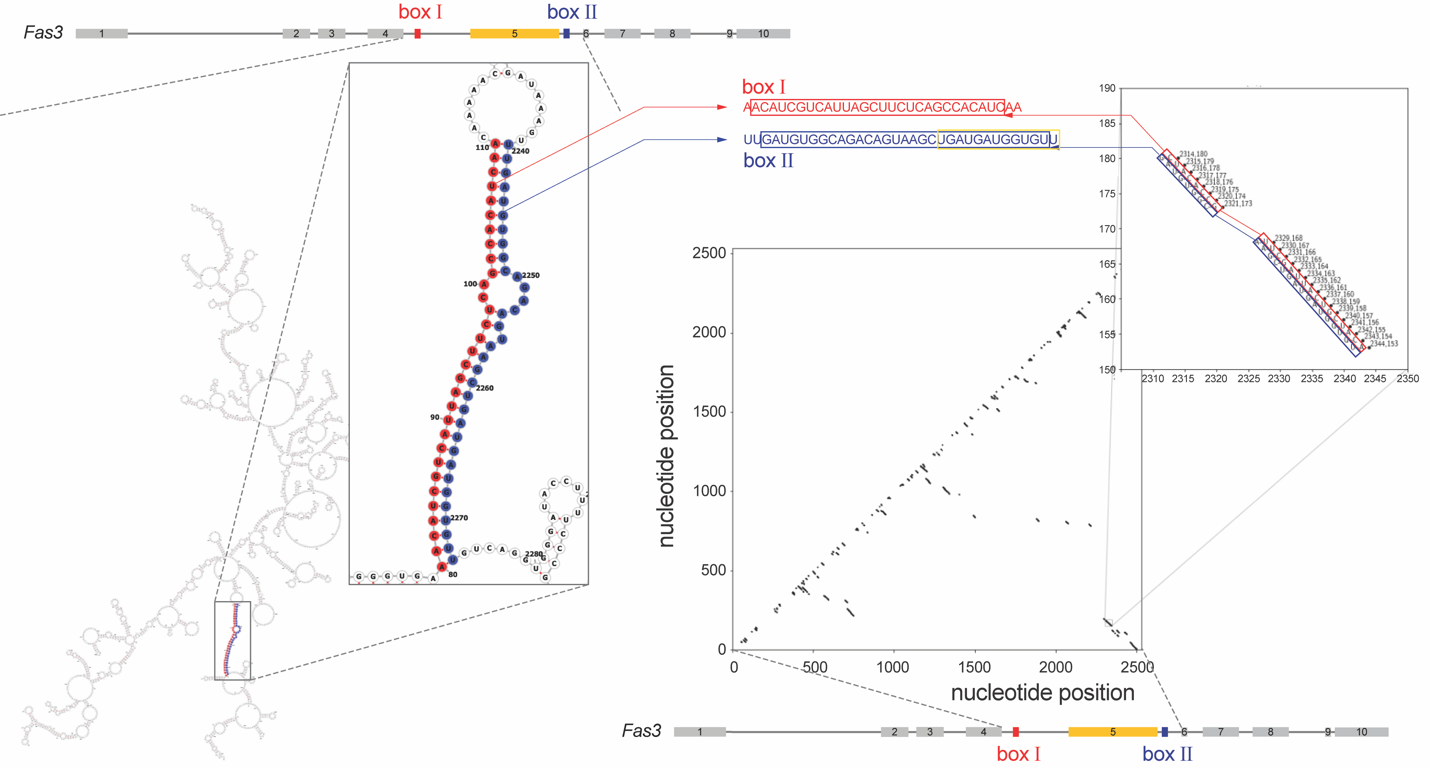

**Figure S1. Determination of splice relevant boxes in the mRNA of *Fas3*.** The matrix shows the nucleotide binding probabilities (P > 0.8) of the sequence of the *Fas3* mRNA that captures the exon 5 and flanking introns. The inset points to a stem formed from box sequences that are near splice sites (red/blue boxes), overlapping with complementary sequences (yellow box). To refine these boxes, the energetically most stable RNA structure of the underlying mRNA subsequence confirms the box sequences through the emergence of a stem in the underlying secondary structure, providing final box sequences.

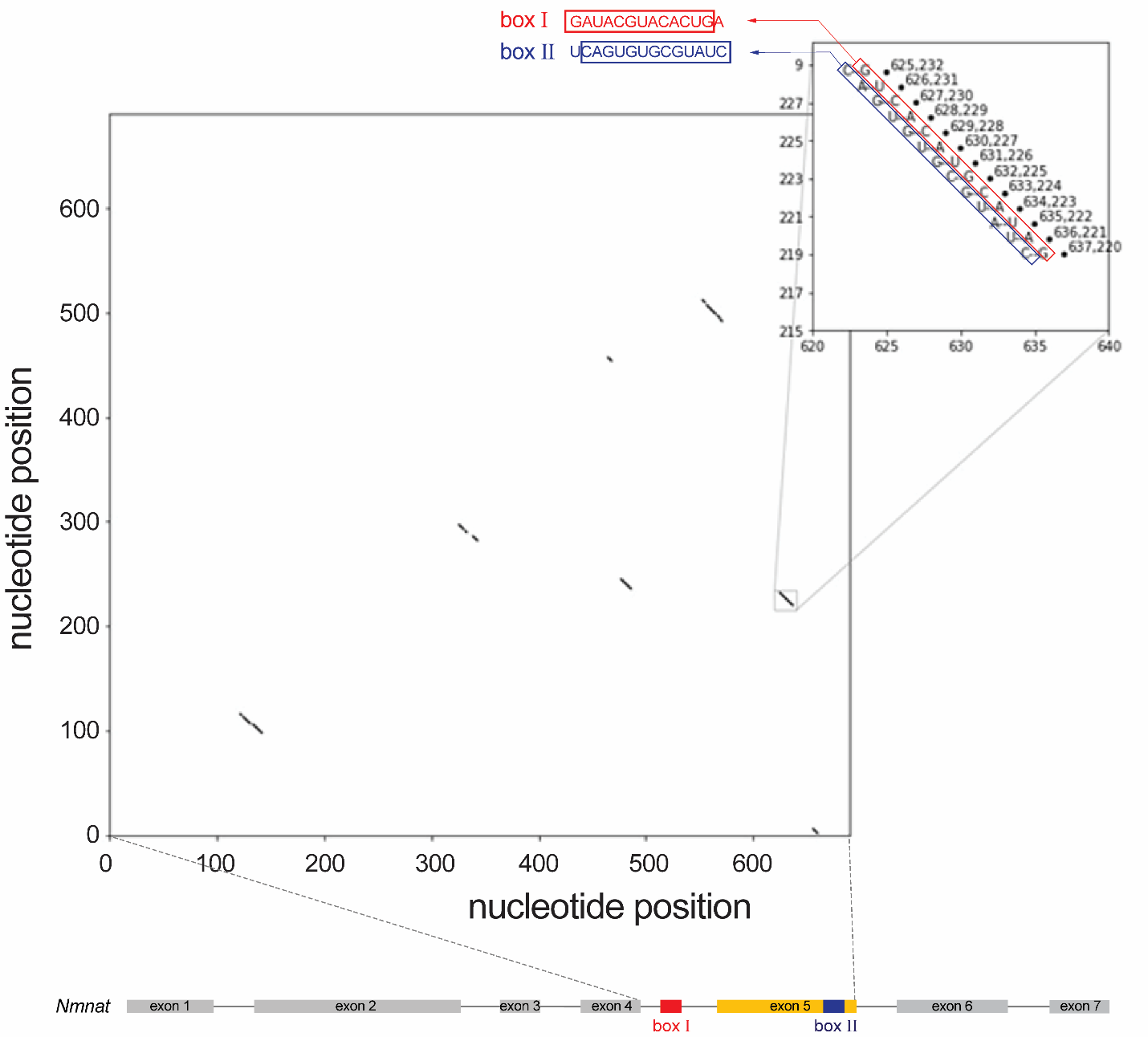

**Figure S2. Determination of splice-relevant boxes in the mRNA of *Nmnat*.** The matrix indicates nucleotide binding probabilities (>0.8) in the subsequence of the pre-mRNA of *Nmnat*, that captures exon 5 and flanking introns. A stem forms from sequences near splice sites (boxes around nucleotides) that overlap with previously found complimentary box sequences.

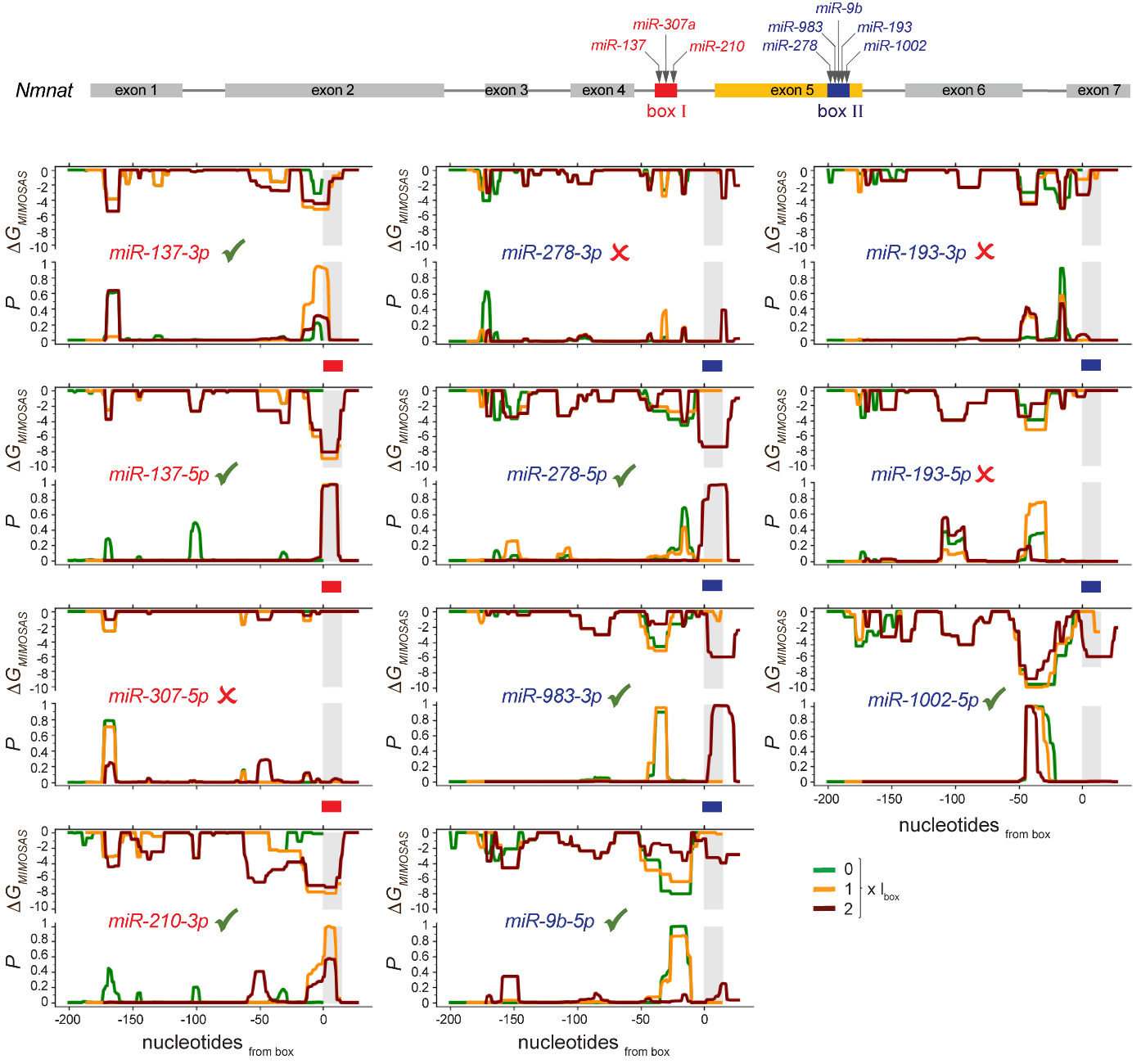

**Figure S3. Determination of microRNA candidates for *Nmnat* splicing.** Focusing on box I and II, we considered a sequence window of multiples of corresponding box lengths around a given box on the *Nmnat* mRNA and calculated *ΔG_MIMOSAS_* as well as the probability that microRNA candidates indeed bind a given box.

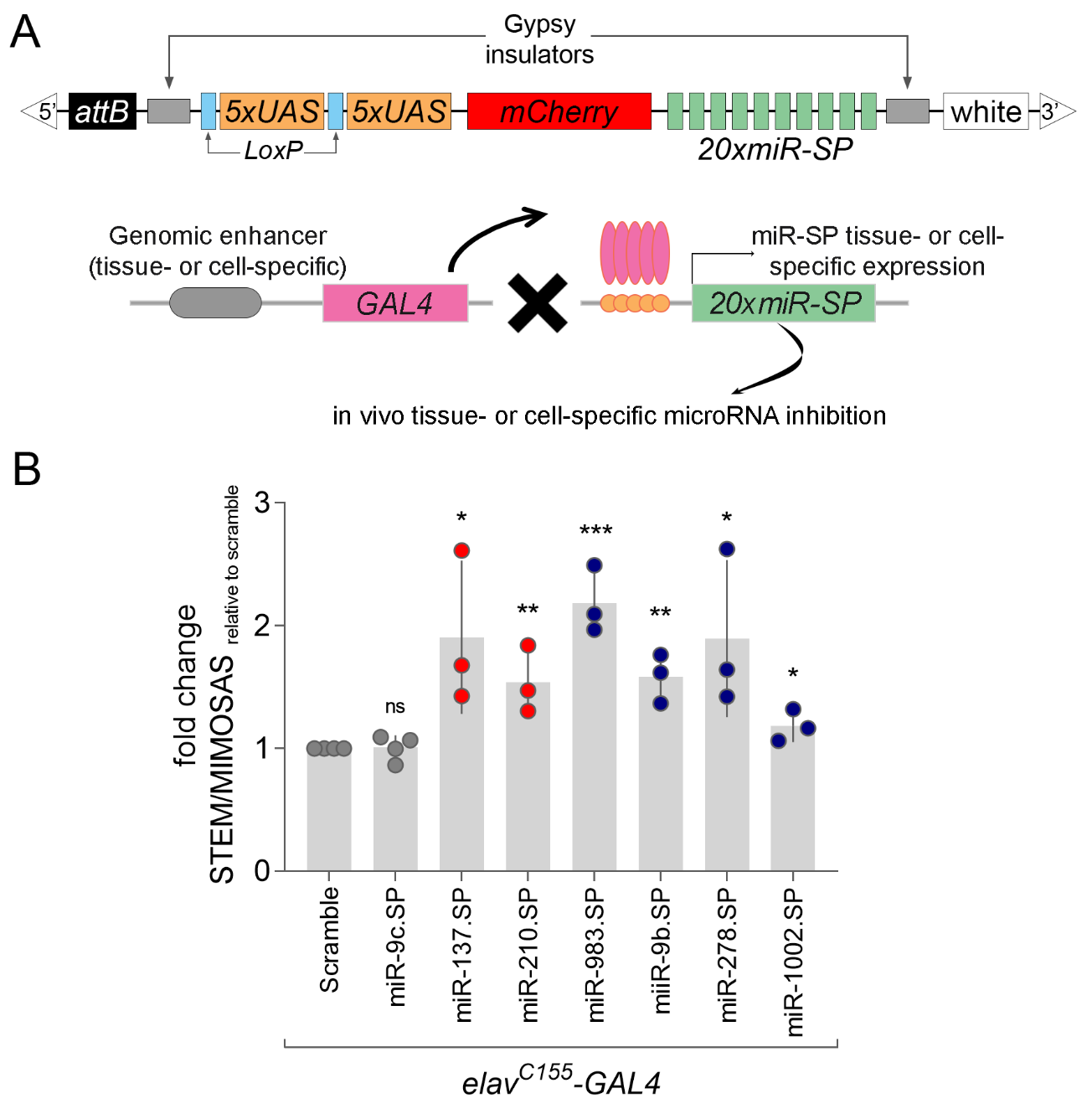

**Figure S4.** **Knockdown of the endogenous microRNAs by miRNA sponges affects MIMOSAS outcome.** (**A**) A gene structure diagram of the microRNA sponge transgenic fly line. 20 microRNA binding sites (green) with mismatches at positions 9-12 were inserted downstream of mCherry in a UAS-containing attB vector. The resulting transgenic animals can be crossed to specific Gal4 lines to achieve a tissue- or cell-specific expression. **(B)** The scatter plot of the ratio between the endogenous RA (STEM) and RB (MIMOSAS) of *Drosophila Nmnat* measured by qRT-PCR in the brains overexpressing miRNA sponges by pan-neuronal driver *elav^C155^-GAL4*. The ratio of RA vs RB in the scramble sponge expressing group was set to 1, and the fold changes were displayed. All data were presented as mean±s.d. ***P≤0.001, **P≤0.01, *P≤0.05, unpaired Student’s *t* test; n≥3, triplicate sampling.

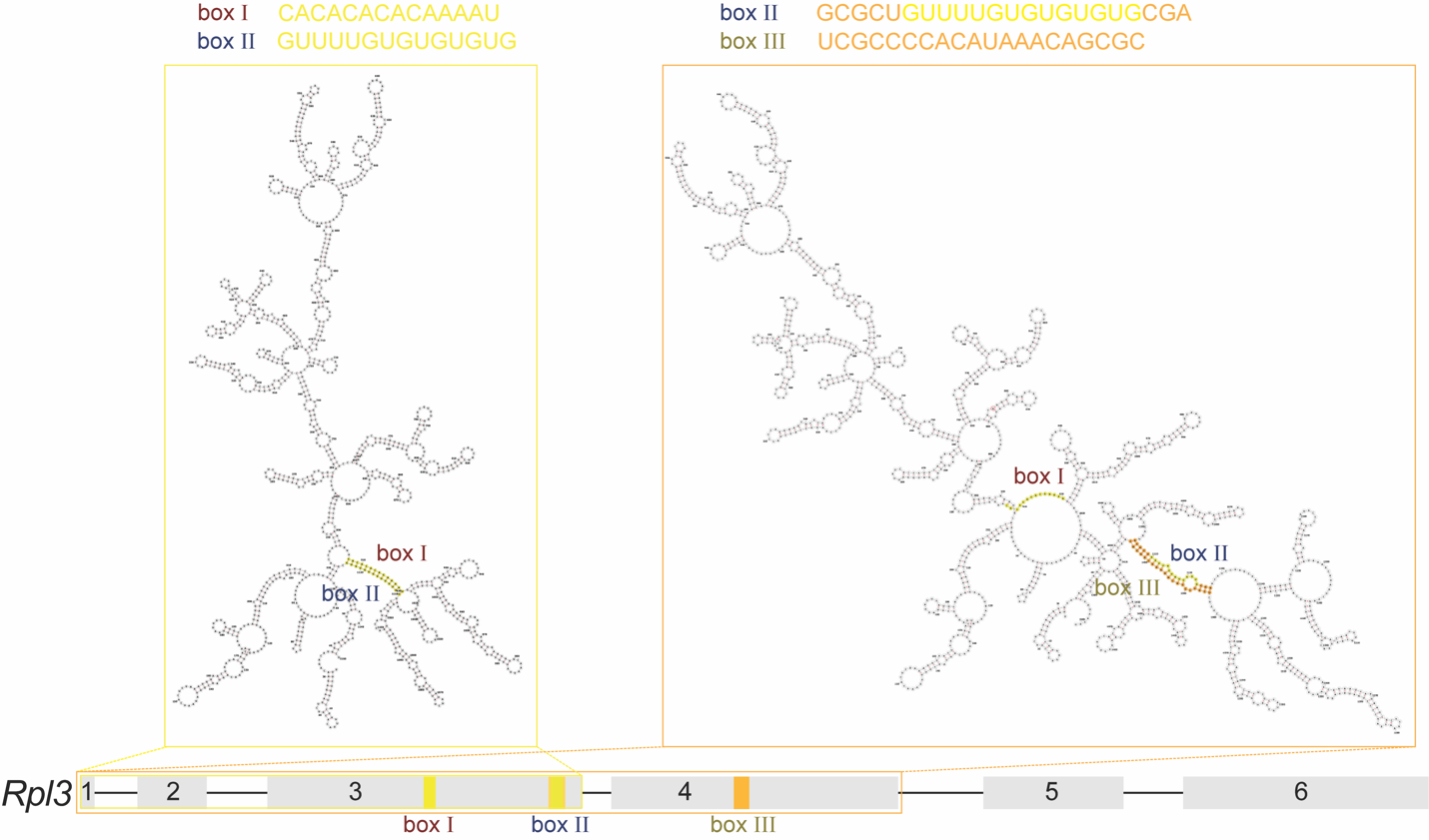

**Figure S5. Determination of nested boxes in *Rpl3*.** Considering the subsequence of *Rpl3* that captures exon 1, 2 and 3 (yellow box), the most stable RNA secondary structure features a formation of a set between box I and II. Focusing on the subsequence of *Rpl3* that captures exon 1, 2, 3 and 4 (orange box), the stem between box I and II dissolved while box II formed a wider stem with box III.

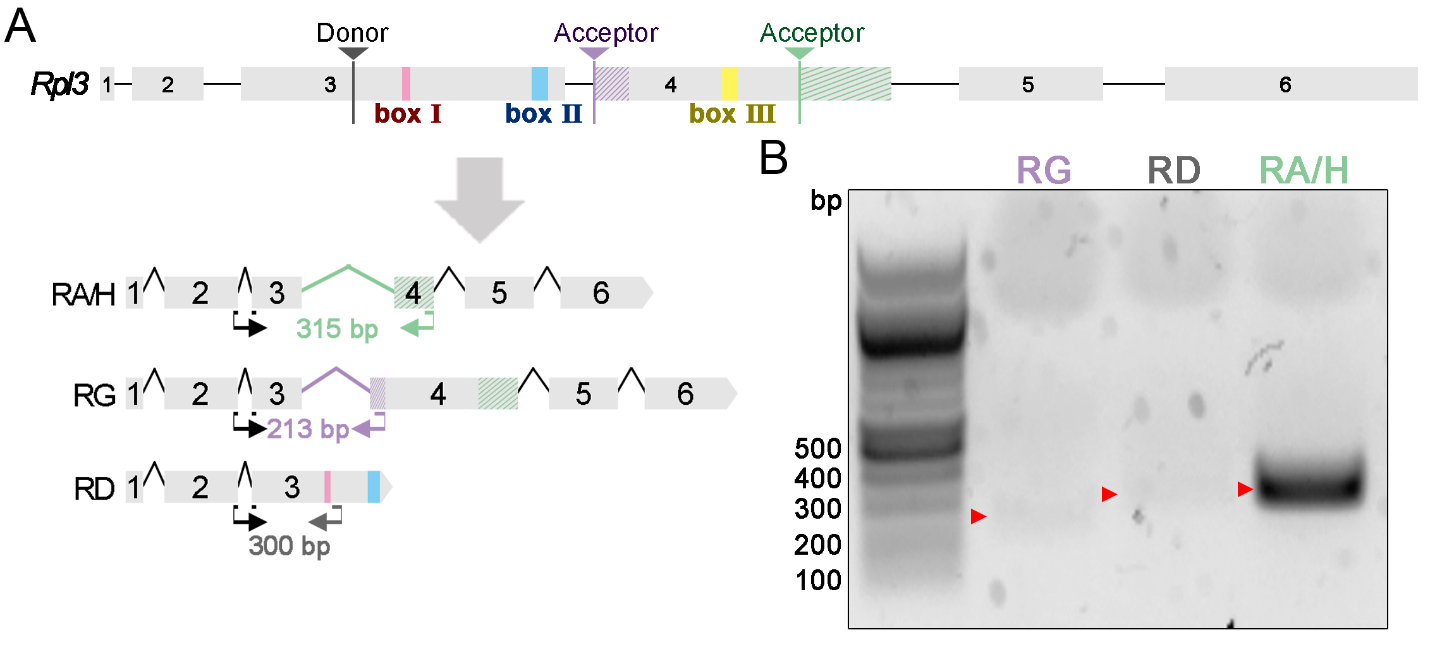

**Figure S6.** ***Drosophila Rpl3* is alternatively spliced into various mRNA variants.** (**A**) Diagram of *Rpl3* gene structure and predicted spliced mRNA variants. Box I (red), II (blue) and III (yellow) mark the sequences that form the stem-loop structures that are required for alternative splicing. Black arrows indicate the sequence location of shared forward primer (F-Rpl3-E2/3). Green (R-Rpl3-RA/H), purple (R-Rpl3-RG) and grey (R-Rpl3-RD) arrows are reverse primers marking the sequence locations for each specific mRNA variant. **(B)** RT–PCR of total RNA from head extracts of wild-type flies (*yw*) using three primer sets as indicated in **(A)**.

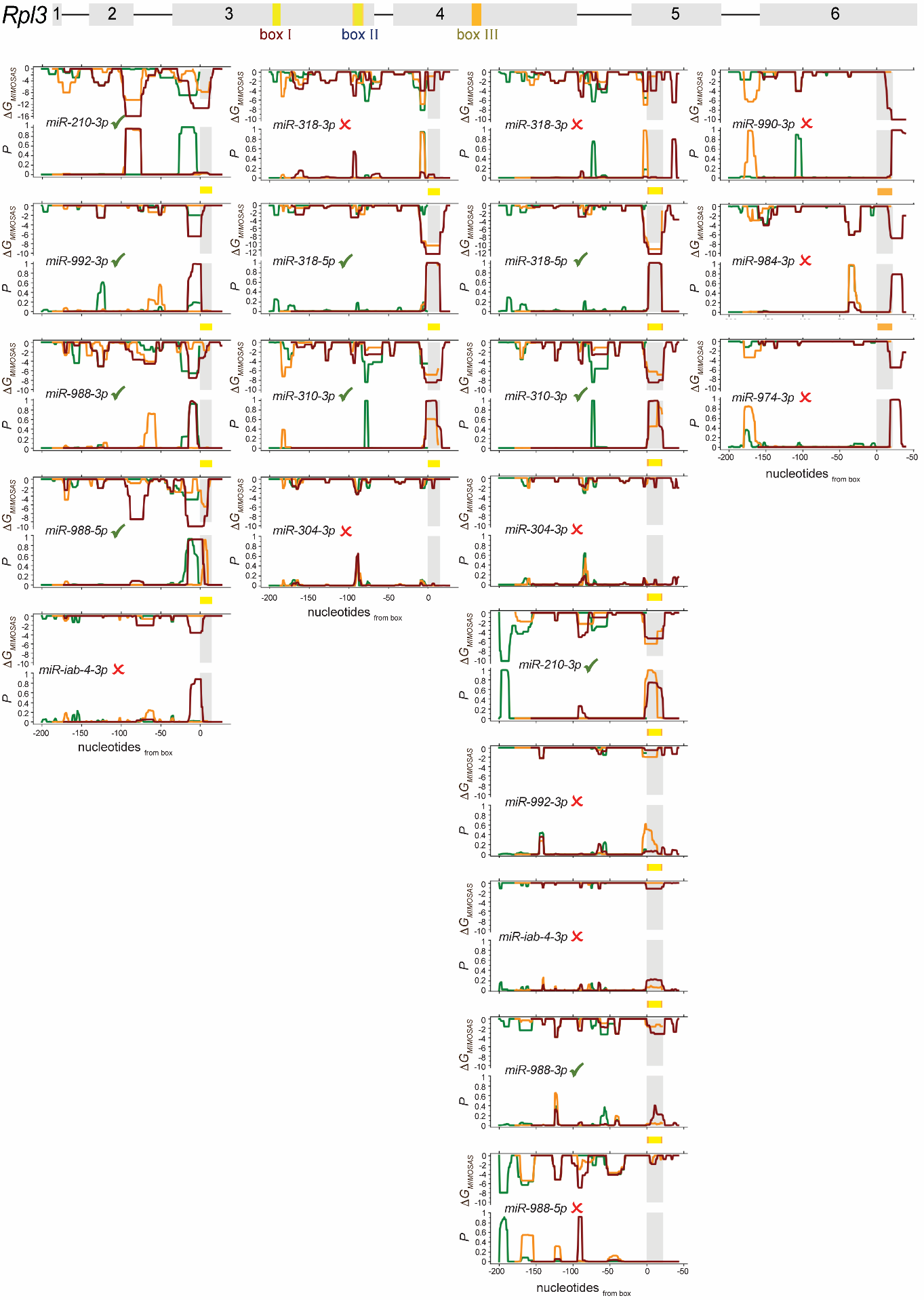

**Figure S7. Determination of microRNAs binding boxes I, II and III in *Rpl3*.** Further investigating the efficacy of miRs to interfere with splice-relevant secondary structures of the underlying Rpl3 mRNA through box I, II and III we considered sequence windows of multiples of corresponding box lengths around a given box and calculated *ΔG_MIMOSAS_* as well as the probability that a miR indeed binds a given box.

**Table S1. A list of primers used in this study.**

| **mRNA detecting** | |
| --- | --- |
| **F-Rpl3-E2/3 R-Rpl3-RD**  **R-Rpl3-RG** | 5’- CCTGGCTCCAAGATCAACAAG-3’  5’- CGTTTCATCATCCAGACTCGAG-3’  5’-GCTCACACAACGTTTAGCGATTTG-3’ |
| **R-Rpl3-RA/H** | 5’-CTGCGAGTGGGCAATCACAC-3’ |
| **Pre-microRNA cloning** | |
| **F-9c**  **R-9c** | 5’- AATTCATTTTTGCTGTTTCTTTGGTATTCTAGCTGTAGATTGTTTCACGCACATTGTATATCATCTAAAGCTTTTATACCAAAGCTCCAGCTTAAATC-3’  5’- TCGAGATTTAAGCTGGAGCTTTGGTATAAAAGCTTTAGATGATATACAATGTGCGTGAAACAATCTACAGCTAGAATACCAAAGAAACAGCAAAAATG-3’ |
| **F-210**  **R-210** | 5’- AATTCAAAGGTGCTTATTGCAGCTGCTGGCCACTGCACAAGATTAGACTTAAGACTCTTGTGCGTGTGACAGCGGCTATTGTAAGAGGCCATAGAAGCAACAGCCC-3’  5’- TCGAGGGCTGTTGCTTCTATGGCCTCTTACAATAGCCGCTGTCACACGCACAAGAGTCTTAAGTCTAATCTTGTGCAGTGGCCAGCAGCTGCAATAAGCACCTTTG-3’ |
| **F-304**  **R-304** | 5’- AATTCGCAGCATTGAATAATCTCAATTTGTAAATGTGAGCGGTTTAAGCCATTTGACGCACTCACTTTGCAATTGGAGATTGCTCGAGACTGCC-3’  5’- TCGAGGCAGTCTCGAGCAATCTCCAATTGCAAAGTGAGTGCGTCAAATGGCTTAAACCGCTCACATTTACAAATTGAGATTATTCAATGCTGCG-3' |
| **F-988**  **R-988** | 5’- AATTCGACGGCGGTACCGGGCATTTTGGGTGTGTGATTTGTAGCAAAGTGATATGTATTTGATCATCCCCTTGTTGCAAACCTCACGCCAAAGATGATCTGCGAC-3’  5’- TCGAGTCGCAGATCATCTTTGGCGTGAGGTTTGCAACAAGGGGATGATCAAATACATATCACTTTGCTACAAATCACACACCCAAAATGCCCGGTACCGCCGTCG-3’ |
| **F-992**  **R-992** | 5’- AATTCATTTTCCCAAGTGCCTGGTATCAGCAAAGTGTTATTTTTTATGTTTATGTAAAGTACACGTTTCTGGTACTAAGTACTTCGAGAAAGTTACCC-3’  5’- TCGAGGGTAACTTTCTCGAAGTACTTAGTACCAGAAACGTGTACTTTACATAAACATAAAAAATAACACTTTGCTGATACCAGGCACTTGGGAAAATG-3’ |
